# Regional *Plasmodium falciparum* subpopulations and malaria transmission connectivity in Africa detected with an enlarged panel of genome-wide microsatellite loci

**DOI:** 10.1101/2024.03.08.584049

**Authors:** Martha Anita Demba, Edwin Kamau, Jaishree Raman, Karim Mane, Lucas Emenga-Etego, Tobias Apinjo, Deus Isheghoma, Lemu Golassa, Oumou Maiga, Anita Ghansah, Marielle Bouyou-Akotet, William Yavo, Milijoana Randrianarivelojosia, Fadel Muhammadou Diop, Eniyou Oriero, David Jeffries, Umberto D’Alessandro, Abdoulaye Djimde, Alfred Amambua-Ngwa

## Abstract

Unravelling the genetic diversity of *Plasmodium falciparum* malaria parasite provides critical information on how populations are affected by interventions and the environment, especially the evolution of molecular markers associated with parasite fitness and adaptation to drugs and vaccines. This study expands previous studies based on small sets of microsatellite loci, which often showed limited substructure in African populations of *P. falciparum*. Combining several short tandem repeat detection algorithms, we genotyped and analysed 2329 polymorphic microsatellite loci from next-generation sequences of 992 low-complexity P*. falciparum* isolates from 15 sub-Saharan African countries. Based on pairwise relatedness, we identified seven subpopulations and gene flow between the Central and Eastern African populations. The most divergent subpopulation was from Ethiopia, while unexpected unique subpopulations from Gabon and Malawi were resolved. Isolates from the Democratic Republic of Congo shared ancestry with multiple regional populations, suggesting a possible founder population of P. falciparum from the Congo basin, where there was stronger geneflow eastwards to Tanzania, and Kenya. and Malawi. The most differentiated microsatellite loci were those around the *P. falciparum* dihydropteroate synthase (*Pfdhps*) gene associated with sulphadoxine resistance. Haplotypes around the *Pfdhps* gene separated the West, Central, and East Africa parasite populations into distinct clusters, suggesting independent local evolution of *Pfdhps*-associated sulphadoxine resistance alleles in each African region. Overall, this study presents genome-wide microsatellites as markers for resolving P. falciparum population diversity, structure, and evolution in populations like Africa, where there is high gene flow.

## Introduction

Malaria remains one of the most important parasitic diseases, particularly in sub-Saharan Africa where five countries (Nigeria, the Democratic Republic of Congo (DR-Congo), Uganda, Mozambique, and Angola) account for just over 50% of the global case burden, with many of these countries also significantly contributing to malaria-related deaths (WHO 2022). Although progress has been made towards effectively controlling malaria, the situation remains unstable, with resurgence of cases in areas of extremely low transmission, potentially fuelled by high human movements across the continent (McAuliffe 2021), drug and insecticide resistance or changes in the climate. The high-resolution mapping of local parasite diversity, and the identification of the origin of infections and signatures of selection or differentiation could help guide interventions strategies and for assessing the impact of their implementation on the evolution of parasite and vector populations.

Determining the local parasite population structure and the origin of infections requires user friendly tools and good quality baseline data. This has often employed several sets of nuclear and non-nuclear single nucleotide polymorphism (SNP) panels for *Plasmodium falciparum* (Daniels et al. 2008; Campino et al. 2011; Preston et al. 2014). A recent genome-wide SNPs analysis characterised the African *P. falciparum* populations into several regional subpopulations (west, central, east and the horn-of-Africa)(Amambua-Ngwa et al. 2019). Microsatellites, especially a set of 12 loci developed over two decades ago, have also been widely used to determine population structure and the relatedness of infections (Anderson et al. 2000). The increasing availability of different *P. falciparum* genome sequences has facilitated the development of even larger microsatellite sets (Liu et al. 2020; Han et al. 2022). These new microsatellite panels have proven useful in resolving the degree of connectivity between different parasite populations at a more granular scale and generating information that can be exploited to interrupt parasite flow between major sources and sinks (Tessema et al. 2019).

Microsatellites can also be used to detect and resolve the history of signatures of positive selections, some of which are associated with drug resistance (Mallick et al. 2013). The evolution of such signatures can also be tracked over time as shown for the large region in *P. falciparum* chromosome 6 known to code for an amino acid transporter (PfAAT1:) that enhances chloroquine resistance (Amambua-Ngwa et al. 2016; Amambua-Ngwa et al. 2023). In addition, microsatellite characterisation across the whole parasite genome is now possible which can be used to refine global parasite population structure, diversity, connectivity, transmission, and other population genetic analyses(Han et al. 2022).

The analysis presented here focuses on a subset of African *P. falciparum* populations collected through the Pathogens genomic Diversity Network Africa (PDNA) (Ghansah et al. 2014) and reported in the MalariaGEN Pf6 data release (MalariaGen et al. 2021). Microsatellite diversity, population structure and connectivity across Africa *P. falciparum* geopolitical populations were determined. Using a variety of approaches, a subset of 2329 microsatellite loci provided increased resolution of *P. falciparum* malaria parasite population structure across Africa. We detected significant gene flow between the central and eastern African parasite populations and strong differentiation around the dihydropteroate synthase (*Pfdhps)* gene associated with resistance to the antimalarial, Sulphadoxine.

## Materials and methods

### Populations

Sequences used in this study were generated from *P. falciparum* isolates collected in 15 PDNA countries as part of a larger MalariaGEN consortium dataset (Pf6 data release) (MalariaGen et al. 2021). The 1240 initially selected isolates were likely monoclonal (i.e. being monogenomic as determined from an inbreeding coefficient (Fws)>0.9). The isolates represented five African sub-populations previously identified by SNP analysis: West (The Gambia, Senegal, Nigeria, Ghana, Cote d’Ivoire, Mali and Guinea), Central (Cameroon, Gabon), South-Central (Democratic Republic of Congo) and East (Kenya and Tanzania), Southeast (Malawi and Madagascar) and the Horn of Africa (Ethiopia), (Supplemental_Table_S1), (Amambua-Ngwa et al. 2019).

### Microsatellite mining

For each isolate, Fastq sequence files were downloaded from the European Nucleotide Archive (ENA) and aligned to the *P. falciparum* 3D7 reference genome sequence (version3, October 2012 release) using the *bwa mem* algorithm. The generated Bam files were filtered to remove unmapped reads, reads with unmapped mates and nonprimary alignment reads, and then sorted. A reference catalogue of short tandem repeat regions in the 3D7 reference genome was created using tandem repeat finder (TRF) ^2^ with the following parameters: match = 2, mismatch = 7, indel = 7, minimum alignment score = 10, and maximum period size = 6 to identify microsatellites. The output was converted into bed and data.region index files for the genome-wide microsatellite mining with lobSTR (Gymrek et al. 2012) and RepeatSeq(Highnam et al. 2013) algorithms, respectively. Homopolymers and repeat regions with more than one insertion were removed from the output variant call format (VCF) files, and the resulting microsatellite VCF merged and sorted. To minimise the percentage of missingness, an iteration removing loci at low call rates and then samples with high genotype missingness was repeated from >=50% to <10% in steps of 5%. A custom script was used to find consensus microsatellite loci with similar motifs, position, and number of repeats from lobSTR and RepeatSeq. Retaining not more than 10% missingness for samples or loci resulted in a final dataset of 992 samples and 2329 loci for lobSTR; and 844 samples and 189 loci for *RepeatSeq* (Supplemental_Table_S2). Both datasets were merged, and duplicates removed (Supplemental_Table_S2). Based on the median genomic position across the repeat region, each locus was classified as intronic or non-coding. Repeat motifs were also classified as complex or simple, where complex motifs could be ambiguously defined (e.g., AT or TA) or interrupted by other polymorphisms, while simple motifs were non-ambiguous and included non-AT nucleotides.

For the 992 isolates retained by lobSTR and RepeatSeq, we explored GangSTR(Mousavi et al. 2019) for maximum likelihood tandem repeat length calls. Following filtration, 8776 loci were retained with a missingness of less than 20% (Supplemental_Table_S3). From these, there were 5765 polymorphic loci, which we analysed in parallel with the merged data from lobSTR and RepeatSeq, but only the latter was retained as the main data for this report.

### Population diversity and complexity

Genome-wide heterozygosity was measured for each locus per parasite subpopulation using adegenet and poppr packages in R. For each isolate, the complexity of infection was re-estimated using a modified inbreeding coefficient, *F_ws,_ computed using the formula *F_WS_ = 1−(H_W_/H_S_), where H_W_ is the within-individual heterozygosity and Hs is the within-population heterozygosity(Roh et al. 2019). H_W_ was calculated using the formula H_W_ = 1-(mean(1/N_A_)^2^) where N_A_ is the number of alternative alleles. Hs was calculated using the formula H_S_=1-(((AD_R_/(AD_R_+AD_A_))^2^ + ((AD_A_/(AD_R_ + AD_A_))^2^) where AD_R_ is the allelic depth for the reference allele and AD_A_ is the allelic depth for the alternative allele.

Multi locus linkage disequilibrium (LD) was calculated as index of association aggregated across all loci with poppr or for each pair of loci using a customised R script employing the LDKl function (which calculates LD statistics for multiallelic markers) found in the genetic analysis package (gap) in R. LD express as D′ was tested for significance using the chi-square test.

### Population structure and admixture

Loci within coding regions, non-coding regions, or those with motifs other than AT dinucleotide were tested separately for population structure using adegenet in R. For each dataset, the provesti distance was calculated and applied to Principal Coordinate Analysis (PCoA) resulting in 15 coordinates which were further subjected to data reduction with Uniform Manifold Approximation and Projection (UMAP)(Leland McInnes 2018). The hclust function in the R package heirfstat was also applied to UMAP coordinates and the results converted to a newick file format for visualization as a neighbor-joining tree with ITOL. In a third approach, the number of predicted ancestries was first estimated using find.cluster without prior population allocation and the number of clusters from Bayesian Information Criterion (BIC) set at k=6, followed by DAPC data reduction. DAPC coordinates were also further reduced with UMAP and visualised as ggplot scatter plots. A similar approach was applied to GangSTR data, retaining the same estimated k.

To determine the level of admixture between the sub-populations, the SNMF function in the Lea package in R was used to model individual ancestry coefficients and ancestral frequencies. Following the cross-entropy criterion from primary iteration of 10 runs per value of k (from k=2 to k=15), k=6 was selected for later analysis. SNMF qmatrices were further evaluated and visualised as customised bar plots with R pophelper package.

### Population differentiation and isolation by distance (ibd)

The significance of population structure was tested using analysis of molecular variance (AMOVA) with poppr, while differentiation between population pairs was evaluated with Nei’s genetic distance (D) across all loci and per locus using the hierfstat and Bayscan R packages. The relationship between genetic and geographic distance between all pairs of populations was tested using *ibd* (isolation by distance) with significance assessed with the Mantel test (mantel.randtest) in ade4 R package, following 1,000,000 permutations.

### Putative connectivity between parasite populations

Relatedness between each pair of isolates within and between populations was first determined by identity-by-state (IBS), calculated with a custom R code, using the formula; IBS=sum (lengths of similar alleles shared between sample1 and sample 2)/(number of samples - missing loci). The generated pairwise isolate IBS matrix was visualised using a heatmap. The level of connectivity (migration) between population pairs was determined as average IBS for all pairs between both populations and displayed on the map of Africa. Additionally, pairwise relatedness between isolates and populations was estimated as identity-by-descent (IBD), calculated from dominant non-reference alleles at loci with multiallelic calls per sample. The pseudo-phased genotype data was diploidised and used with fastIBD in BEAGLE to determine pairwise IBD. For each pair of samples, the mean genome wide IBD across 10 runs was calculated. We determined highly related pairs that may be because of recent transmission or common recent ancestry, by introducing isolates from geographically distal Asia as a control population, where significant recent transmission or ancestry with Africa is not expected. For this analysis, microsatellites from 539 monoclonal samples from 6 Asian countries (Bangladesh, Laos, Cambodia, Thailand, Vietnam, Myanmar) identified by lobSTR, were merged with the African dataset and common loci retained for IBD calculations as above. The overlap in IBD distribution between Asian and African samples was considered as the null (ancestral) distribution, not determined by recent relationships that may be due to linked transmission events. Highly related isolates were defined as parasite pairs with IBD at 3 standard deviations beyond the mean (or beyond the 95% quantile) of the distribution of IBD between isolate pairs from Africa. The resulting highly related IBD pairs were visualised in a connectivity map linking the sites of sampling. Additionally, pairwise relatedness within populations was also assessed using the basic allelic sharing index (Blouin’s Mxy) in demerelate package in R.

## Results

### Genome-wide polymorphic microsatellite loci are common across the genome

Most of the 428,471 potential short tandem repeat segments in the reference genome consisted of mononucleotide and complex short tandem repeat motifs. Polymorphic repeat motifs genotyped from deep sequence *reads* ranged from mononucleotide to hexanucleotide (six). Excluding homopolymer runs, the initial genome-wide dataset included 1208 samples and 141,822 and 65,500 polymorphic microsatellite loci from lobSTR and RepeatSeq, respectively. LobSTR identified twice as many loci compared to RepeatSeq (Supplemental_Fig_S1). Following merging and filtration down to overall missingness of 10%, a final dataset of 992 samples and 2*329* microsatellite loci with 2 to 24 alleles were retained (Supplemental_Table_S2). For GangSTR, 8776 polymorphic loci were *initially* genotyped in 992 samples. 5666 loci with 2 to 30 alleles, dominated (69%) by “AT/TA” dinucleotide motifs, were polymorphic and ha*d* less than 10%missingness. They were used *only* for population structure and per locus differentiation analyses (Supplemental_Table_S3).

The 2329 loci extracted with lobSTR-RepeatSeq were polymorphic in all populations and distributed across all chromosomes in the *P. falciparum* 3D7 reference genome, with more loci identified for larger chromosomes (Supplemental_Fig_S2). Of the 144 polymorphic microsatellite motifs detected (Supplemental_Table_S4), those with less than 2 nucleotides were more prevalent than longer ones (62.4% di-nucleotides; 19.3% trinucleotides; 11.5% tetra-nucleotide; 4.2% penta-nucleotide; and 2.7% hexanucleotide (Figure S3A). Motifs with only adenine (A) and thymine (T) bases with two to five nucleotides were dominant, while those with guanine (G) and cytosine (C) bases were detected in only 14.86% of the motifs and mostly in complex ones. Most (>60%) of these polymorphic microsatellite loci genotyped and retained were located within coding regions of the genome (Supplemental_Fig_S4).

### Genetic diversity

Most samples and populations had a high microsatellite heterozygosity (multiple alleles called per locus), with a median Nei’s unbiased expected heterozygosity of 0.53 (Supplemental_Table_S5). Heterozygosity was uneven across the genome, with similar patterns in the different country and regional populations (Supplemental_Fig_S5a). Heterozygosity decayed around regions coding for drug resistance loci (PfMDR1, PfCRT, PfDHFR-TS and PfPPPK-DHPS), although this was mostly skewed upstream of downstream the focal gene due to the low density of markers genotyped around their coding regions (Supplemental_Fig_S5b). Overall, the complexity of infections per country as determined by the variant of the inbreeding coefficient, *F_WS,_ was substantially higher compared to that derived and reported from SNP data(Amambua-Ngwa et al. 2019). The mean per sample *F_WS_ was 0.7 and ranged from 0.69 – 1.00 (figure 1a). Isolates from Gabon had the highest

**Figure 1:**
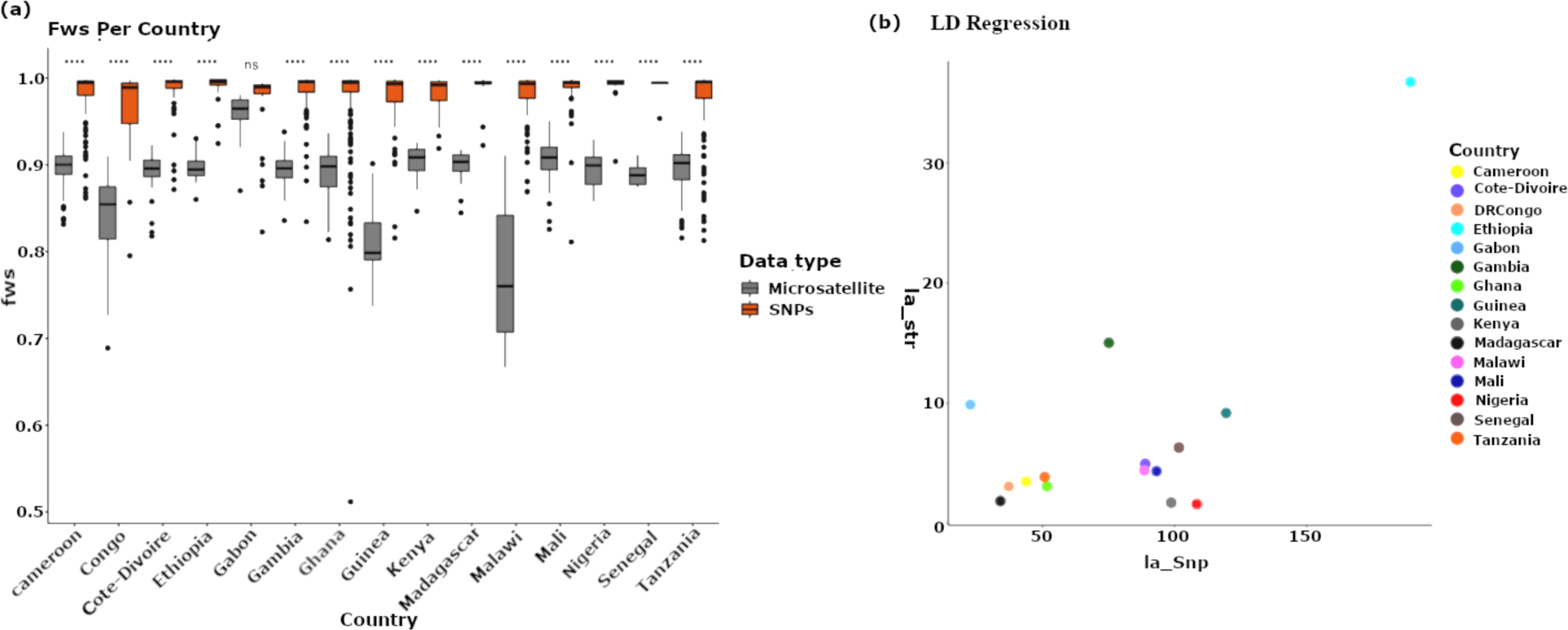
Complexity of infection determined by inbreeding coefficient *Fws for isolates from PDNA countries in Africa. a) Barplot of the distribution of *Fws within countries calculated with SNP and microsatellite data. b) Correlation of LD (index of association) determined from SNP and microsatellite data per population.

*F_WS_ values, while those from Malawi had the lowest.

The overall population LD determined by index of association (Ia) between loci showed Ethiopian isolates having the highest LD followed by isolates from The Gambia (Supplemental_Table_S4). LD determined by microsatellites was much lower but mostly had the same extent per population compared to that determine from SNPs, especially for East African populations (figure 1b).

### Distinct sub-populations of P. falciparum in Africa

Based on cumulative variance explained by BIC from initial PCoA and find.clusters - DAPC analyses, the isolates from the 15 African countries were distributed into about 7 genetic clusters (Supplemental_Fig_S6a and S6b), captured mostly by the first 3 principal components of DAPC (Supplemental_Fig_S6b). A scatterplot of axes 1-3 distinguished these 7 subpopulations (figure 2a and Supplemental_Fig_S6c). These groups roughly aligned with the geographic source of the isolates from west, central and eastern Africa. Surprisingly, the first three components of DAPC also separated Central African isolates from Gabon and Cameroon, as well as Malawian subgroup from the rest of East Africa and DR-Congo. Combining either DAPC or PCoA with UMAP further resolved Malawian isolates into two groupings, one specific to Malawi and a smaller subset of isolates clustering with those from Madagascar and Eastern Africa (Kenya and Tanzania) (Figure S6d). These unique country populations were also evident from the IBS-based NJ tree figure 2b). This substructure was maintained when only loci in exons were considered (Supplemental_Fig_S6e), but not with loci from introns which grouped mostly into a single cluster, except for isolates from Ethiopia and Malawi (Supplemental_Fig_S6f). Distinct subpopulations and the split within Malawi and between Cameroon and Gabon remained evident after removing loci with dinucleotide AT motifs (Figures S6g) or based on the IBS only (Figure S6h). Considering only loci with Nei’s Fst>=0.1 between geographic populations, this resulted in multiple genetic clusters with high degree of geographic heterogeneity, except for isolates from Ethiopia and Gabon (Figure S6 I & J). Across all different clustering and projection approaches, isolates from DR-Congo dominated a cluster that included isolates from all other regions. The sub-structure revealed by lobSTR-RepeatSeq data was not evident with the parallel data from GangSTR, except for the unique Ethiopian population. The subclusters from Malawi and Gabon were only evident when PCoA-UMAP was applied to GangSTR genotypes (Supplemental_Fig_S7).

**Figure 2.**
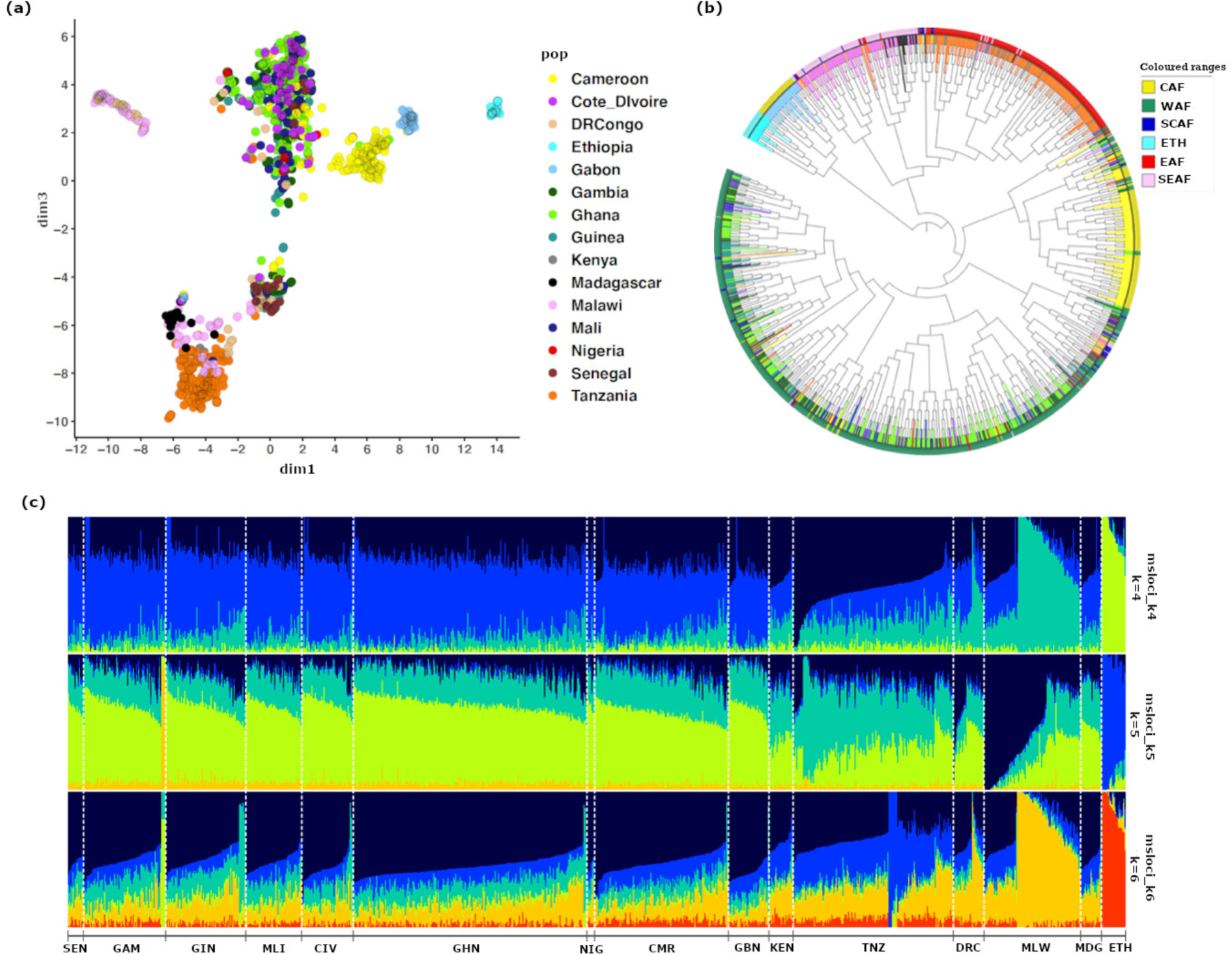
Population structure and ancestry of the *P. falciparum* isolates from Africa. a) Scatter plot axes 2 and 3 of UMAP clustering of combined coordinates from find cluster, and DAPC, b) neighbour joining tree of isolates annotated by region of origin. c) Admixture-like bar plots of modelled ancestry proportions for 4 to 6 ancestries shown in rows with isolates groups and sorted by country of origin at the bottom (from left to right, SEN=Senegal, GAM=Gambia, GIN=Guinea, MLI=Mali, CIV=Cote D’Ivoire, GHN=Ghana, NIG=Nigeria, CMR=Cameroon, GBN=Gabon, KEN=Kenya, TNZ=Tanzania, DR-Congo=Democratic Republic of Congo, MLW=Malawi, MDG=Madagascar, ETH=Ethiopia).

Cross-entropy analysis of likely ancestries maximised at an estimated k ranging from 4 - 6 (Supplemental_Fig_S8). Modelling ancestry proportions from these 4 – 6 backgrounds, all populations were predominantly admixed, except for the more unique population in Ethiopia (figure 2c). Common patterns were observed for isolates belonging to the main West, Central, and East African geographic groupings. Two levels of admixture were seen for Malawi, with a smaller sub-group showing patterns similar to Madagascar and East African isolates while the dominant unique Malawian sub-group remained distinct.

### Pairwise relatedness and putative connectivity between country populations

Genetically related isolates based on IBD and IBS were detected within and between populations. Overall, pairwise relatedness was stronger for isolates sampled within than between country and regional populations (figure 3a). Isolates from Ethiopia and Malawi had the highest within population relatedness by IBS (figure 3b) and IBD (Supplemental_Fig_S9), with two the subgroups in Malawi evident from distinct pairwise ancestries. Considering pairs of isolates beyond the 95^th^ percentile of genome-wide relatedness, highly related pairs were more evident across borders within regions. In West Africa, a relatively high proportion of related infections were observed between Ghana and Guinea, while the strongest cross border relatedness connected mostly DR-Congo with Eastern Africa, especially Malawi and Tanzania (figure 3c). There was also a relatively higher percentage of highly related pairs within East African countries compared to other regions, with Ethiopia and Malawi having the highest proportion of within-population related pairs (figure 3d).

**Figure 3.**
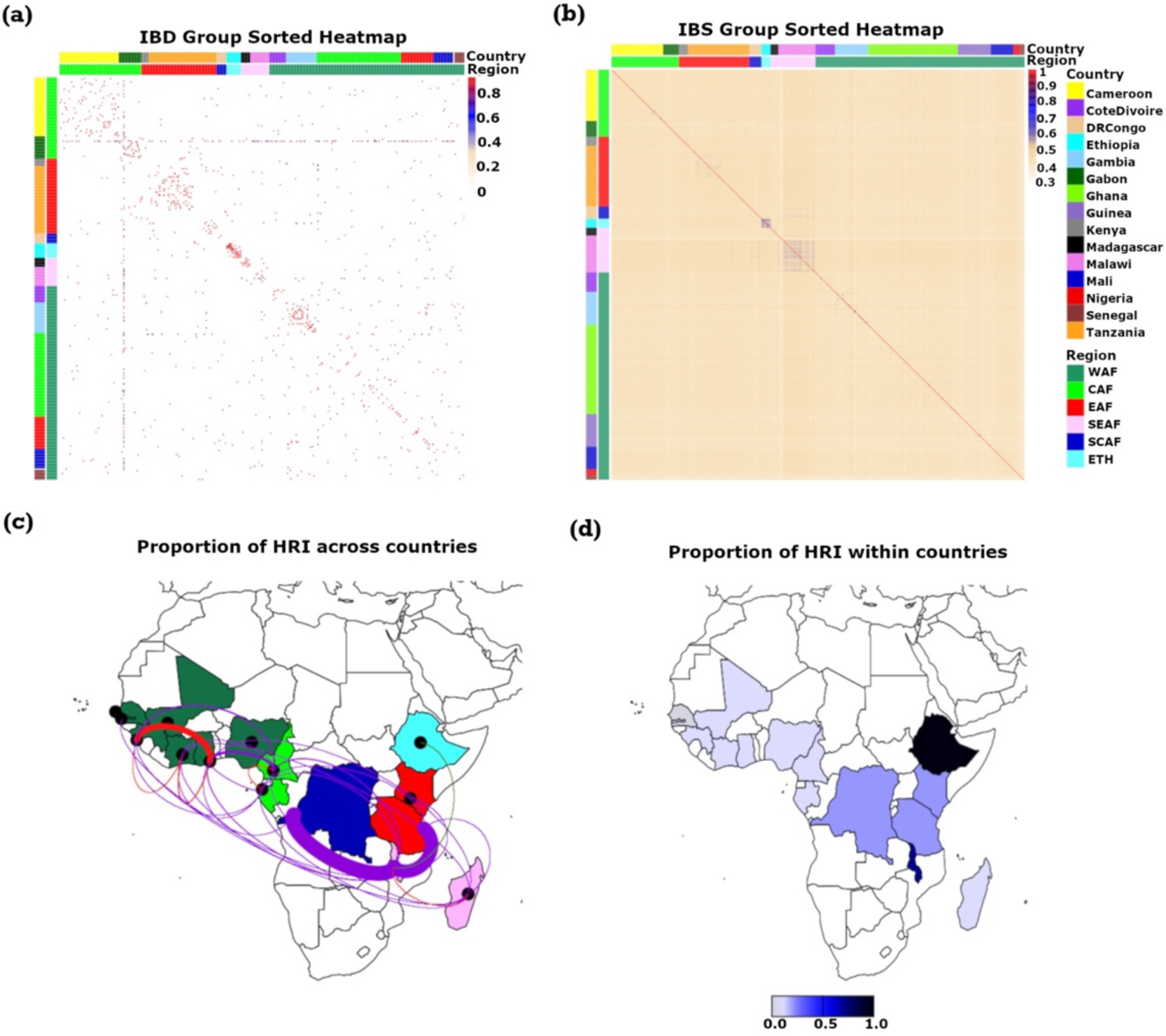
IBD and IBS between and within populations. a) heatmap of pairwise IBD transformed as - log_10_ of IBD, b) heatmap of pairwise IBS, c) Connectivity between country populations determined by proportion of high related pairs (HRI), with thickness of lines proportional to the proportion of HRI connecting infections within regions (red), between different regions (purple), and with Ethiopia (green). Regional population are colour coded as dark green for west Africa (WAF), yellow green for central Africa (CAF), blue for southcentral Africa (SCAF), red for east Africa (EAF), cyan for Ethiopia (ETH) and pink for Southeast Africa (SEAF). d) Within population IBS density, scaled from light to dark blue from low to high IBS respectively.

### Pairwise population differentiation

Overall, pairwise population differentiation by Nei’s genetic distance was generally low between all pairs of populations, with Ethiopian population being most distant from all other country populations. Fst was therefore non-significant for most population pairs, and AMOVA showed similar variance within populations compared to between populations, phi = 0.016. Clustering of pairwise population genetic distances identified Gabon, Malawi, Ethiopia as unique from the East African (DR-Congo, Tanzania and Kenya), West-Central African (Mali, Ghana, Cote d’Ivoire, and Nigeria), and West African west coast grouping that included Gambia, Guinea, Senegal, but surprisingly Cameroon (figure 4a). Mentel’s tests for isolation by distance (*ibd)* that compares Nei’s genetic distance against geographic distance was non-significant (Supplemental_Fig_S10). However, there was a trend of increasing genetic distances by physical distance when isolates from Ethiopia and Gabon were excluded (figure 4b).

**Figure 4.**
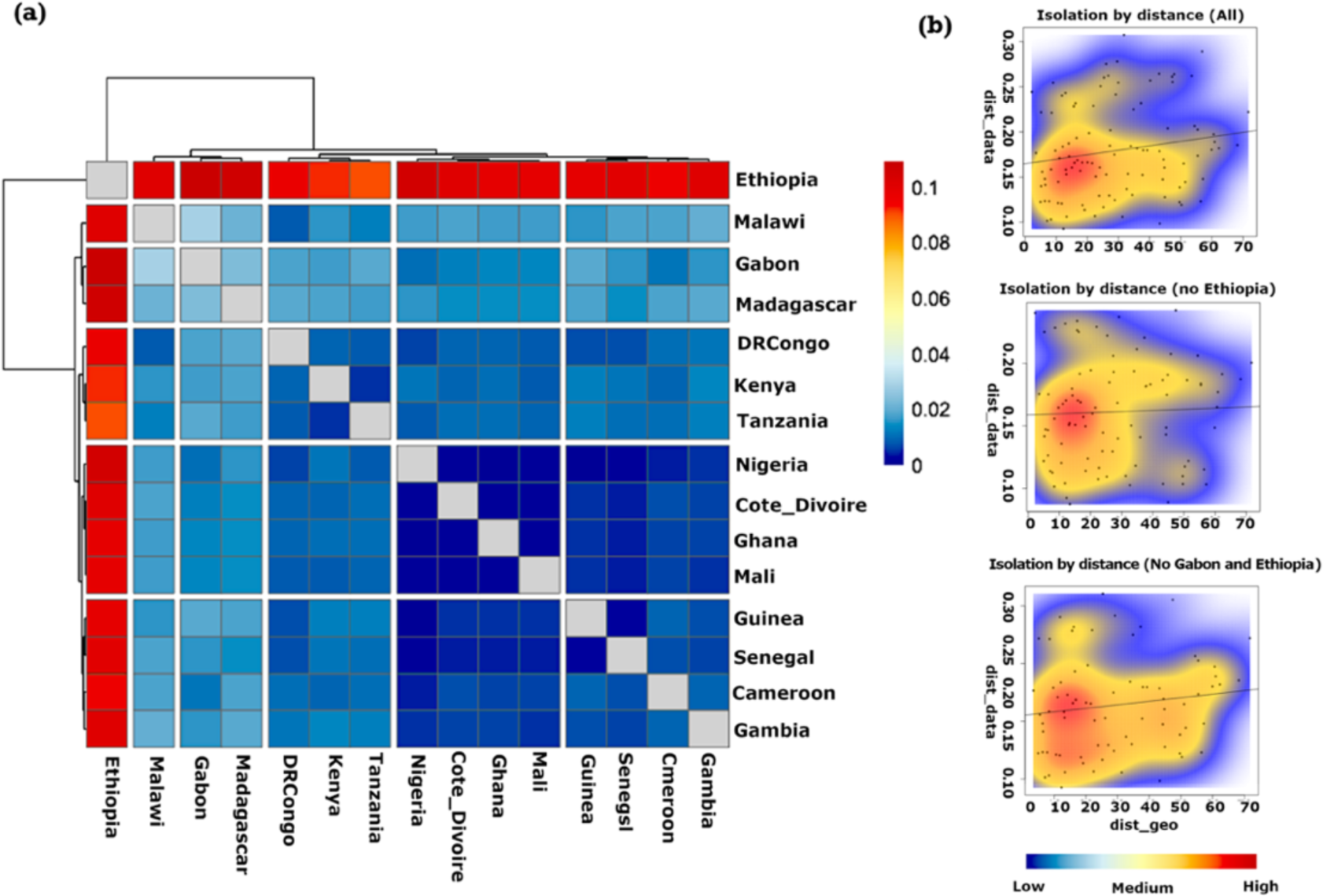
Population clusters from high Nei’s pairwise population Fst repeat loci and isolation by distance showing, a) Clustered heatmap of pairwise Nei’s Fst between *P. falciparum* in African populations, and b) Contour plots of isolation by distance (*ibs*) Mentel’s test of geographic (dist-geo) against genetic (dist-data) distances. The line shows the correlation between the two distance matrices. The first row of column b shows *ibs* for all 15 African countries. The second row is without Ethiopia and the third is when isolates from Gabon and Ethiopia are excluded.

### Microsatellite loci differentiation

Between all populations, 196 microsatellites extracted with lobSTR-RepeatSeq had an Dest >0.025 and were thus considered as differentiating populations (Supplemental_Table_S6). Differentiation indices were higher for GangSTR data, with 166 loci having Dest >0.05. Combined, these differentiating loci included those in 234 gene coding regions, loci in 37 genes detected as highly differentiating by data extracted by all methods (Supplemental_Table_S7 and S8). Both sets of data showed the highest density of differentiating loci on chromosome 8, around the coding region of dihydropteroate synthetase (*Pfdhps*) gene (figure 5a). Clustering infections using only loci around the *Pfdhps* gene grouped infections into three major clusters, i.e., West Africa, East Africa, and infections from DR-Congo and Gabon, with Cameroon linking the West and Central Africa blocks. Isolates from Madagascar were mostly unique (Figure 5b). The most distant populations for this genomic region were those from West Africa against Kenya and Ethiopia.

**Figure 5.**
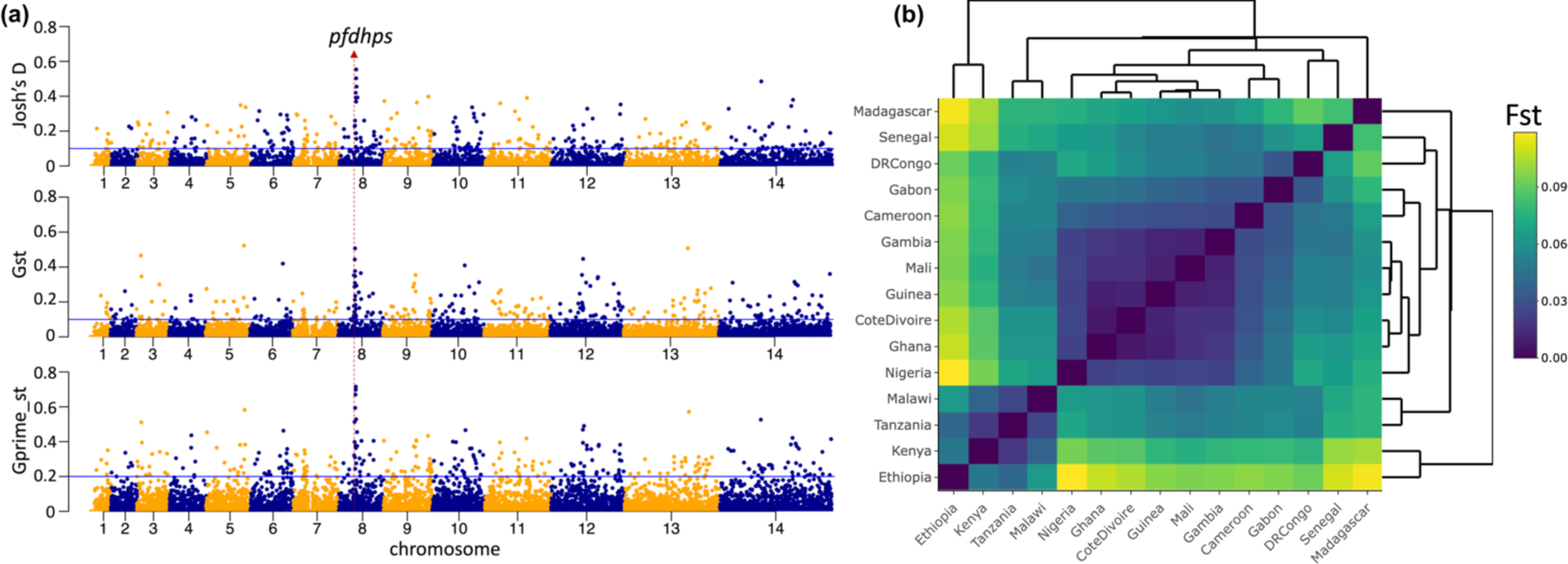
Differentiating microsatellite between African populations, showing strong differentiation at the DHPS locus. a) Manhattan plots of differentiation indices per locus presenting Jost’s D, Gst and Gprime_St from top to bottom panel. b) Cluster heatmap of infections from African populations based on haplotypes around DHPS gene.

## Discussion

With the development of new tools for microsatellite repeat mining from deep sequence data, this study explored open-access *P. falciparum* whole genome sequences generated from collaboration between mostly PDNA consortium and the MalariaGEN community project to further assess *P. falciparum* diversity in Africa (MalariaGen et al. 2021). The subset of highly heterozygous microsatellites was extracted with lobSTR and RepeatSeq, which had previously been shown to enable high-quality genotyping of repeat alleles for population genetic studies of positive selection(Amambua-Ngwa et al. 2016). This included highly diverse genome-wide loci from 992 low complexity infections across 15 countries, allowing increased resolution of the population structure, relatedness and recent patterns of selection and movements of this important malaria parasites in Africa (Schaid et al. 2004). The data also enabled the detection of higher levels of infection complexity, attributable to the higher allelic diversity of microsatellites compared to bi-allelic SNP data(Amambua-Ngwa et al. 2019; MalariaGen et al. 2023). Due to higher rates of transmission, complex polygenomic malaria parasite infections are common in Africa, although this was not the case for infections from Ethiopia, which had also been shown previously with SNPs to be the least complex but most divergent from the rest of continent (Amambua-Ngwa et al. 2019). This is not surprising as malaria transmission in Ethiopia is extremely low, with highly clonal infections and significant geographic barriers to spatial gene flow with the rest of Africa (Tessema et al. 2020; Abera et al. 2021; Tadele et al. 2022). Consequently, isolates from Ethiopia also had relatively higher index of association, a measure of LD, probably driven by the higher inbreeding and lower infection complexity (Tadele et al. 2022). The overall LD, which is a function of the level of recombination, also correlated with those determined by SNPs, further confirming the usefulness of this set of microsatellites loci for population genetic studies of *P. falciparum* in Africa.

By combining the set of microsatellites with additional clustering and dimension reduction approaches, African *P. falciparum* isolates were distinguished into seven genetic subclusters beyond the six regional population clusters shown by SNP analysis (Amambua-Ngwa et al. 2019). However, like with SNP based studies, population structure was predominantly defined by region or country of origin of the isolates. Irrespective, the set extracted here allowed for a higher resolution of population structure than previously described for any set of genetic variants (Anderson et al. 2000), including microsatellite genotypes from HipSTR (Han et al. 2022) and those from GangSTR from this study. This increase in resolution was highlighted by isolates from Gabon clustering away from those from Cameroon, although these Central African countries share a long border. It is probable that the river and dense tropical forest separating the two countries limits free gene flow between the parasite populations, against perceived human migration fuelled by trade. In addition, the isolates analysed from Cameroon were mostly collected around the Mount Cameroon, hundreds of kilometres from the Gabonese border, and therefore unlikely to have significant geneflow and recent relatedness with Gabon. This result is in contrast with data from SNPs that grouped the same Cameroonian and Gabonese isolates together into a single central African cluster (Amambua-Ngwa et al. 2019). It is therefore a demonstration of the additional power of microsatellites to resolve population structure even at high geographic proximities (Fola et al. 2020). This is also unlike described for *P. vivax*, where SNPs provided a better resolution (Fola et al. 2020), maybe because of a low AT content, high SNP variation and higher population substructure compared to *P. falciparum* (Carlton et al. 2008; Rougeron et al. 2020). A second highlight was the clustering of isolates from Malawi into two distinct groups, one of which was unique to Malawi and the other related to East African isolates. The isolates from Malawi were collected from Chikwawa and Zomba districts in the south of the country bordered by Mozambique(Ocholla et al. 2014). These results could be due to differences in the collection period, although we note that the districts are separated by rivers and Blantyre, the country’s capital. Malawi is a land-locked country, where migration of populations from different countries could introduce diverse lineages. This was supported by the significant IBS and IBD between Malawi, Congolese, and Tanzanian populations.

Congolese isolates dominated a single cluster in population structure analysis that included members from all other regions but shared the highest relatedness with East Africa. This suggest that *P. falciparum* isolates from this region may have played a central role in the ancestry or the recent expansion of *P. falciparum* across Africa. Our findings together with the previous SNP-based chromosome painting analysis hypothetically support possible zoonotic historical spill over of malaria parasites into human around the Congo basin, from the great apes population either in Cameroon or the Congos (Krief et al. 2010; Liu et al. 2010)(Sibley 2019). The gorilla population in African which carry the ancestral *P. parafalciparum* malaria parasite inhabit the grand Congo basin and equatorial forest of Central Africa that spans from Western Cameroon, DR-Congo to Rwanda and Uganda in East Africa. Humans may have been first exposed to *P. falciparum* in this geographical area with the parasite diverging from its ancestor as humans migrated (Duval et al. 2010; Prugnolle et al. 2011; Loy et al. 2017). Expanded genome wide analysis of human and primate malaria in the Congo basin could shade more light on malaria history.

Although malaria parasites expanded and became endemic in different and distal regions of Africa, the populations remained connected as shown by poor differentiation (non-significant isolation by distance). This mixing was most obvious in West Africa, were populations were less distinct and pairwise genetic relatedness was the weakest due to the intense transmission and consequent frequent recombination of diverse parasite genotypes (WHO 2022). Complex infections with mixed genotypes are also common in West Africa. This was in contrasts with East Africa where highly related infections within and across borders were more common, especially in Malawi, where control interventions had been scaled up (Cohee et al. 2022), with resultant reduction in transmission intensity and infection complexity. The population from Malawi and Tanzania were strongly connected to DR-Congo where political instability may be promoting significant recent human migration along with their infection to the East. Relative to the West, East African isolates also showed a stronger sub-structure that was at odds with observed high geneflow. This pattern is like what has been observed in southeast Asia, where infections are less complex and population substructure is evident even with genetic connectivity (Shetty et al. 2019).

Based on pairwise per locus differentiation, the genomic region around the PfDHPS gene was the most divergent. Mutations in this gene are associated with Sulfadoxine resistance. Microsatellite variation around the gene grouped isolates into regional subclusters suggestive of unique haplotypes in regional populations. It has been postulated and partially shown with SNP and other microsatellite analysis that the PfDHPS locus may have evolved differently in different regions (Vinayak et al. 2010; Alifrangis et al. 2014). Haplotypes circulating in Eastern and Western Africa are different (Pearce et al. 2009), although the most common resistant haplotypes were introduced from Southeast Asia into East Africa and subsequently selected following the use of sulphadoxine-pyrimethamine (SP) for malarial treatment. It is also possible that the selection and spread of other PfDHPS SP-resistant haplotypes in Africa occurred prior to the introduction of SP, as antifolate drugs such as cotrimoxazole were already being widely used across malaria endemic countries before SP (Jelinek et al. 1999), (Morjan and Rieseberg 2004). Although microsatellites are generally considered not to be the targets of selection, their proximity to advantageous evolving loci could lead to hitch-hiking and evidence of reduced heterozygosity and LD that is common around selective sweeps^30^. Our results further support the usefulness of microsatellites as a tool for analysing the evolution of malaria parasites, with the possibility of developing new tools for early identification of parasite populations evading control interventions.

The results this study show the utility of microsatellites for analysis of evolution and population dynamics. However, the loci extracted, and data shown are limited to those from PDNA consortium accessible from the open source Pf6 dataset. As most of the specimens were opportunistically collected over several years, this may have significant temporal heterogeneity and they census sample may not be representative of the geopolitical or regional populations named. For some populations, such as Nigeria, the number of isolates were limited and from a single site. Therefore, future studies should be designed for more representative sampling and for development of tools that are suited for haploid and mixed genotyped infections as commonly seen for malaria in Africa. In addition, the validation of some of the population relevant markers identified could enrich panels for targeted genomic surveillance of *P. falciparum*.

In conclusion, genome wide polymorphic microsatellites resolved *P. falciparum* population structure and diversity across sub-Saharan at a higher resolution, identifying unique subpopulations from Gabon, Malawi, and Ethiopia. These loci were also able to detect signatures of selection driven by drug pressure. Overall, the set used here can be further exploited for monitoring populations and informing control programs as elimination strategies are implemented.

## Data Access

The raw sequence reads for *Plasmodium falciparum* malaria parasites included in this study have been published with Pf6 MalariaGEN data release, and accessible via European Nucleotide Archives (ENA). The genotypes used here, and sample IDs are accessible in the supplement.

## Acknowledgement

The authors and PDNA members wish to thank all the patients and guardians who generously agreed to provide blood samples from malaria patients and carriers. We are indebted to all partners of the MalariaGEN Community Project from Africa and Asia, whose data have been included in this analysis. The PDNA. Special thanks to the MalariaGEN team at the Sanger and to Prof Dominic Kwiatkowski (Posthumous) for conceptualizing PDNA and generous data generation through MalariaGEN.

